# Filling of a water-free void explains the allosteric regulation of the β_1_-adrenergic receptor by cholesterol

**DOI:** 10.1101/2021.08.30.457941

**Authors:** Layara Akemi Abiko, Raphael Dias Teixeira, Sylvain Engilberge, Anne Grahl, Stephan Grzesiek

## Abstract

Proteins often contain cavities, which are usually assumed to be water-filled. Recent high-pressure NMR results indicate that the preactive conformation of the β_1_-adrenergic receptor (β_1_AR) contains completely empty cavities (dry voids) of about ∼100 Å^3^ volume, which disappear in the active conformation of the receptor. Here we have localized these cavities by X-ray crystallography on xenon-derivatized β_1_AR crystals. One of the cavities coincides with the binding pocket of cholesterol. Solution NMR data show that addition of the soluble cholesterol analog cholesteryl hemisuccinate (CHS) impedes the formation of the active conformation of the receptor by blocking conserved GPCR microswitches. This wedge-like action explains the function of the cellularly highly abundant cholesterol as a negative allosteric modulator of β_1_AR. The detailed understanding of GPCR regulation by cholesterol via filling of a dry void and the easy scouting for such voids by xenon may provide new routes for the development of allosteric drugs.

## INTRODUCTION

The interior of experimentally determined protein structures often contains cavities, which are not filled by protein atoms. Such cavities are generally thought to be filled by water. However, in some proteins complete voids, not containing any water molecules have been detected by X-ray crystallography or NMR spectroscopy^1–4^. These observations are so far limited to a few model systems, and it is unclear how abundant dry voids are in proteins and to what extent such voids are connected to protein function and structural stability.

Recent high-pressure NMR analysis has shown that such a functional role exists in G protein-coupled receptors (GPCRs)^5^. GPCRs are transmembrane proteins that regulate many vital functions of the human body, and as such are highly attractive drug targets. They recognize a variety of extracellular stimuli and subsequently trigger intracellular downstream signaling cascades^6^. The GPCR function is achieved by the modulation of several conformations encoding inactive and active states, which indicate a high intrinsic flexibility^7–16^. Notably, the static agonist- and antagonist-bound structures are very similar and do not reflect their functional difference^17^. In particular, binary agonist-receptor complexes are in a dynamical equilibrium between a preactive and an active conformation^7^. The active conformation largely corresponds to the conformation in ternary complexes with G protein or G protein-mimicking nanobodies (Nbs) where transmembrane helices (TM) 5 and 6 have moved outward to accommodate the G protein/Nb^18^ and the water-mediated H-bond bridge between Y^5.58^ and Y^7.53^ (YY-lock, superscripts indicate Ballesteros −Weinstein numbering^19^) is closed^7^. In contrast, the preactive conformation of the agonist complex largely resembles the inactive conformation of antagonist complexes with an open YY-lock^7^. G protein recognition of agonist-receptor complexes apparently occurs by conformational selection of the active conformation.

The conformational equilibria of GPCRs are sensitive to many factors. E.g. point mutations, introduced to stabilize the β_1_-adrenergic receptor (β_1_AR) for structural studies, shift the equilibria towards an inactive receptor^8,20–23^. In particular, mutations of the highly conserved^24,25^ Y^5.58^ and Y^7.53^ residues, stabilize β_1_AR by 11 °C, but suppress the formation of the active conformation in binary agonist complexes thereby abrogating G-protein binding^7,8^.

We have recently shown that pressure modulates these conformational equilibria in a mildly thermostabilized β_1_AR mutant (YY-β_1_AR), which contains the Y227^5.58^-Y343^7.53^ motif and is fully capable of G protein binding^5^. When subjected to pressure (midpoint ∼600 bar), YY-β_1_AR is shifted from a mixture of preactive and active conformations to a fully populated active conformation even in the absence of a G protein or G protein-mimicking nanobody. This pressure dependence shows that the active conformation has an about 100 Å^3^ smaller volume than the preactive conformation, which must be due to the collapse of empty (not water-filled) cavities within the receptor-detergent micelle.

An important, extrinsic factor modulating GPCR conformational equilibria are lipids, with cholesterol (CLR) being the most explored^26–30^. CLR is highly abundant in human cells and essential for maintaining cell excitability and homeostasis^30–38^. CLR increases GPCR thermal stability^39–45^ and often acts as an allosteric modulator of GPCR activity^46–51^. The direction of modulation can vary, as CLR e.g. stabilizes the inactive state of CCK1R^47^ and the active state of BLT2R^50^. The molecular mechanisms of such effects are not well understood. They may be caused by direct interactions with receptors or more indirectly by a modulation of membrane physical properties such as fluidity and stiffness.

As CLR is highly insoluble, commonly its more soluble analog, cholesteryl hemisuccinate (CHS) is used in functional and structural studies^44,50^. CHS induces similar thermo-stabilizing effects as CLR^51^ and molecular dynamics simulations show that protonated CHS mimics well many of the membrane modulating properties of CLR^52^. CLR and CHS are the most prevalent lipids resolved in GPCR structures, but often resolution is not high enough to distinguish between both^38,53^. Although the well-defined localization of CLR/CHS in these structures suggests specific binding, no conserved binding sites have been identified^53^, even for closely related GPCRs such as the β_1_- and β_2_-ARs^54– 56^. Instead, CLR or CHS appear at multiple sites, usually in shallow grooves between the transmembrane helices or at the receptor surface^30,53^.

Hydrophobic voids tend to incorporate noble gases^2^. Here we have detected the hydrophobic voids in β_1_AR by crystallography of xenon-derivatized receptor crystals. One of the identified voids colocalizes with the CHS binding site. An NMR analysis of the effect of CHS binding to an agonist-β_1_AR complex shows that CHS restricts the receptor dynamics and hinders the formation of the active state. This is corroborated by a comparison of β_1_AR structures, which reveals that CHS bound to the inactive receptor obstructs the path of subtle, but essential movements required to activate the canonical microswitch GPCR network. Thus CHS apparently acts like a wedge blocking a void necessary for β_1_AR activation. The observed phenomenon of functional dry voids together with the possibility to easily detect such voids by xenon may introduce a valuable new principle to search for relevant drug target sites in proteins.

## RESULTS

### Xenon reveals empty cavities in agonist-bound β_1_AR

To localize the hydrophobic voids within β_1_AR we have obtained crystals of a thermostabilized mutant (TS-β_1_AR)^8^ bound to the agonist isoprenaline and derivatized these with xenon under weak (5–10 bar) pressure. Xenon is easily detected and localized by X-ray crystallography due to its anomalous scattering and high number of electrons^57,58^. As dry xenon immediately destroyed the fragile receptor crystals, the xenon derivatization was carried in a home-built pressurizing device (Figure 1A), where a defined humidity could be set by flowing xenon through a wash bottle before reaching the crystal pressure chamber. An electronic humidity sensor and a glass window in the crystal pressure chamber allowed continuous monitoring of humidity and crystal integrity under a microscope. Best results with no observable crystal damage were achieved by pressurizing the crystals with xenon for 20 min at 5 bar and 100% humidity.

**Figure 1.**
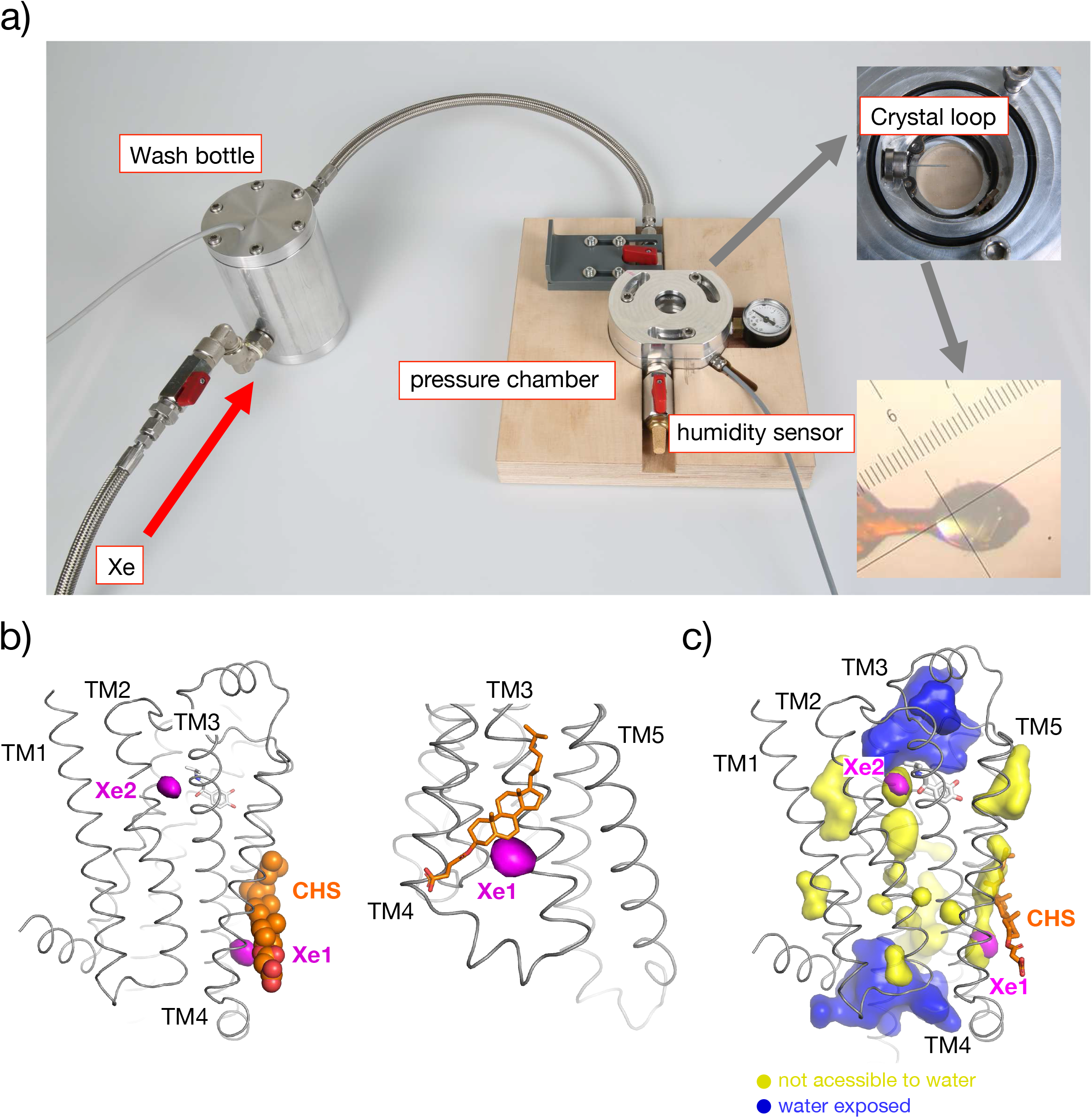
Xenon derivatization of TS-β_1_AR. **a)** Pressure apparatus built for Xe derivatization with humidity control. Pressure chamber, wash bottle, humidity sensor and crystal loop are indicated. Inserts show a close-up of the pressure chamber and a microscopic image of a β_1_AR crystal during Xe pressurization. **b)** Left: Location of the two xenon binding sites (magenta surfaces) identified by anomalous X-ray scattering within the isoprenaline·β_1_AR crystal structure (PDB 2y03). Isoprenaline is represented as gray sticks and CHS as orange spheres. Right: close-up of the Xe1 site, which colocalizes with the CHS molecule (orange sticks) of the crystal structure. **c)** Structure of the isoprenaline·β_1_AR complex (PDB 2y03) and void volumes (shown as surfaces) determined with the program Hollow 1.2.54. Water and lipid molecules were removed before the analysis. Blue/yellow surfaces represent void volumes accessible/inaccessible to water, respectively. Xenon sites Xe1 and Xe2 are shown in magenta and CHS as orange sticks, respectively

The structure of the isoprenaline·β_1_AR complex determined from the xenon-derivatized crystals (Table S1) is almost identical to the previously solved structure of a similar β_1_AR construct in complex with the same agonist (PDB 2y03)^55^ (Figure S1A). Both structures have two receptor molecules in identical arrangement in the asymmetric unit with an overall Cα root-mean-square deviation (RMSD) of 0.705 Å (0.607 Å excluding loops). However, the large anisotropy of the Xe-derivatized crystals prevented localization of the isoprenaline and CHS molecules observed in the 2y03 structure. Two xenon binding sites (Xe1, Xe2) were located unambiguously (signal amplitudes >5σ) within the anomalous scattering Fourier map (Figures 1B,C and S1B, Table S2). Xe1 is positioned towards the intracellular side between TM3 and TM4, and Xe2 towards the extracellular side between TM2 and TM3. Both xenon binding sites constitute crevices, which are formed almost exclusively by hydrophobic side chains (Figure S2A, B).

A computational search for cavities in the 2y03 isoprenaline·β_1_AR structure revealed 15 voids (Figure 1C), two of which are larger and exposed to water at the extracellular and intracellular faces of the receptor. The remaining cavities are smaller and buried in the hydrophobic membrane part of the receptor. Two of these overlap with the Xe1 and Xe2 sites (Figure 1B, C). The smaller void (∼20 Å^3^) largely coincides with the volume of the Xe2 site detected by the xenon anomalous scattering. The second, larger void (∼35 Å^3^) constitutes a hydrophobic groove, which is partially but not completely filled by the CHS molecule in the 2y03 structure (Figure 1C). The detected xenon volume at Xe1 resides in the remaining non-filled part of the groove and is in direct contact with the space occupied by the CHS molecule. Despite not being deeply inserted between the helices, CHS is stabilized by an extensive set of hydrophobic interactions involving TM3, TM4, and TM5 (Figure S2C). Notably, the hydrophilic CHS succinic acid tail is not in direct contact with the receptor, but points towards the solvent thereby leaving the Xe1 part of the hydrophobic groove unoccupied. From this arrangement, it is sterically possible that in the native membrane the sterol moiety of CLR fully occupies the Xe1 pocket.

### CHS obstructs pressure-induced receptor activation

To investigate the effect of CHS on the receptor void volumes, we acquired pressure-dependent ^1^H-^15^N TROSY spectra of the G protein binding-competent ^15^N-valine-labeled isoprenaline·YY-β_1_AR complex^7,8^ in the presence (Figure 2A) and absence (Figure 2B) of CHS. Besides small chemical shift changes and variations in intensity (see below), the valine ^1^H-^15^N resonances are very similar at 1 bar with and without CHS. However, when the pressure is increased to 2500 bar, the spectra diverge strongly. Without CHS, the previously observed^5^ pressure-induced change to the active conformation is evident from strong shifts of the V122^3.33^, V172^4.56^, V202^ECL2^, and V314^6.59^ resonances (Figure 2B) around the orthosteric ligand pocket (see also Figure 3C). In contrast, these active-conformation resonances almost disappear with CHS at 2500 bar. Instead, many resonances are broadened, indicative of an exchange between many conformations (Figure 2A, C). Only few resonances, such as V89^2.52^, V103^2.66^, and V320^ECL3^ remain intense, which must be due to either fast exchange between local subconformations or the absence of local heterogeneity. Monitoring of the protein stability from the intensity of amide ^1^H^N^ resonances revealed that the receptor is strongly destabilized at 2500 bar by the addition of CHS, whereas CHS stabilizes the receptor at 1 bar (Figure S3). Thus CHS apparently blocks the activating motion at 2500 bar, and directs the receptor to a multitude of different unstable conformations. As reaching the active state is strongly hindered by CHS at high pressure, part of the required void volume for activation must comprise the CHS/CLR binding-site.

**Figure 2.**
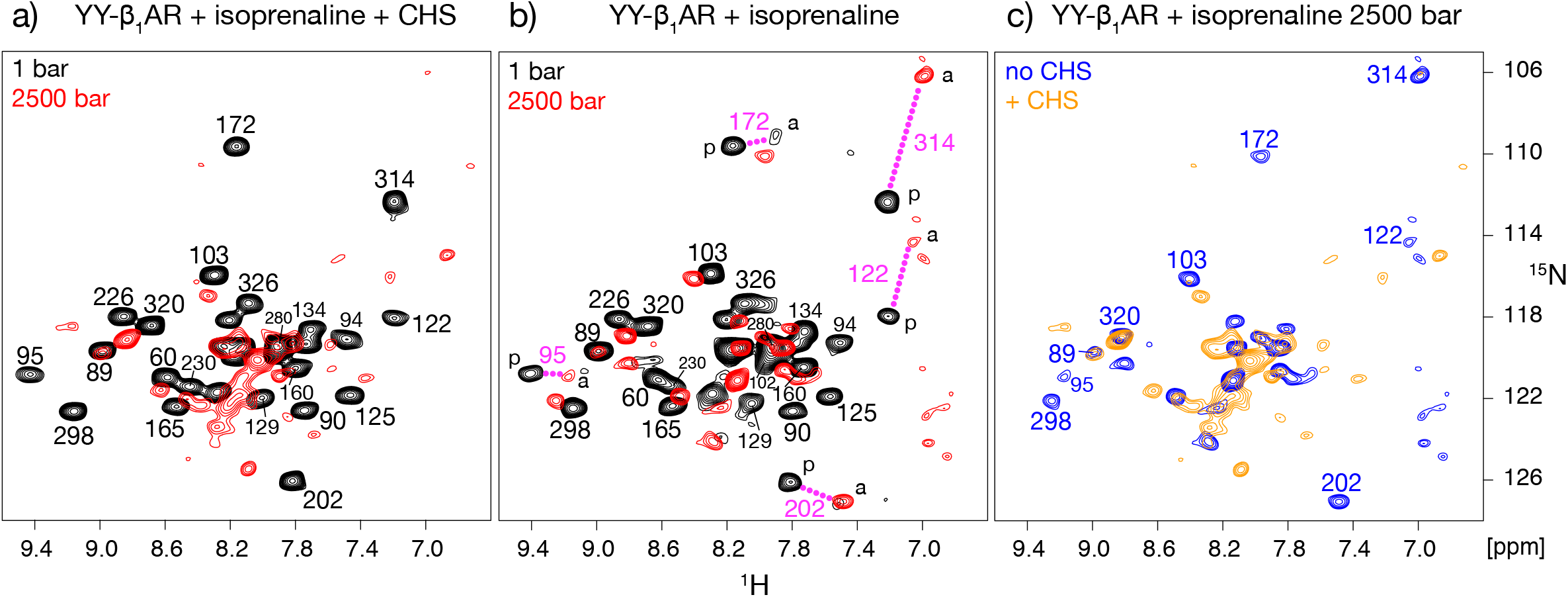
High-pressure NMR analysis of agonist-bound β_1_AR labeled by ^15^N-valine in the presence or absence of CHS. Superposition of ^1^H-^15^N TROSY spectra of **a)** isoprenaline·YY-β_1_AR with 1 mM CHS at 1 bar (black) and 2500 bar (red), **b)** isoprenaline·YY-β_1_AR without CHS at 1 bar (black) and 2500 bar (red), **c)** isoprenaline·YY-β_1_AR with (orange) and without CHS (blue) at 2500 bar. Resonances are marked with assignment information. Resonances connected by a dashed magenta line represent residues with two clearly distinguishable resonances for the preactive (‘p’) and active (‘a’) conformations.

**Figure 3.**
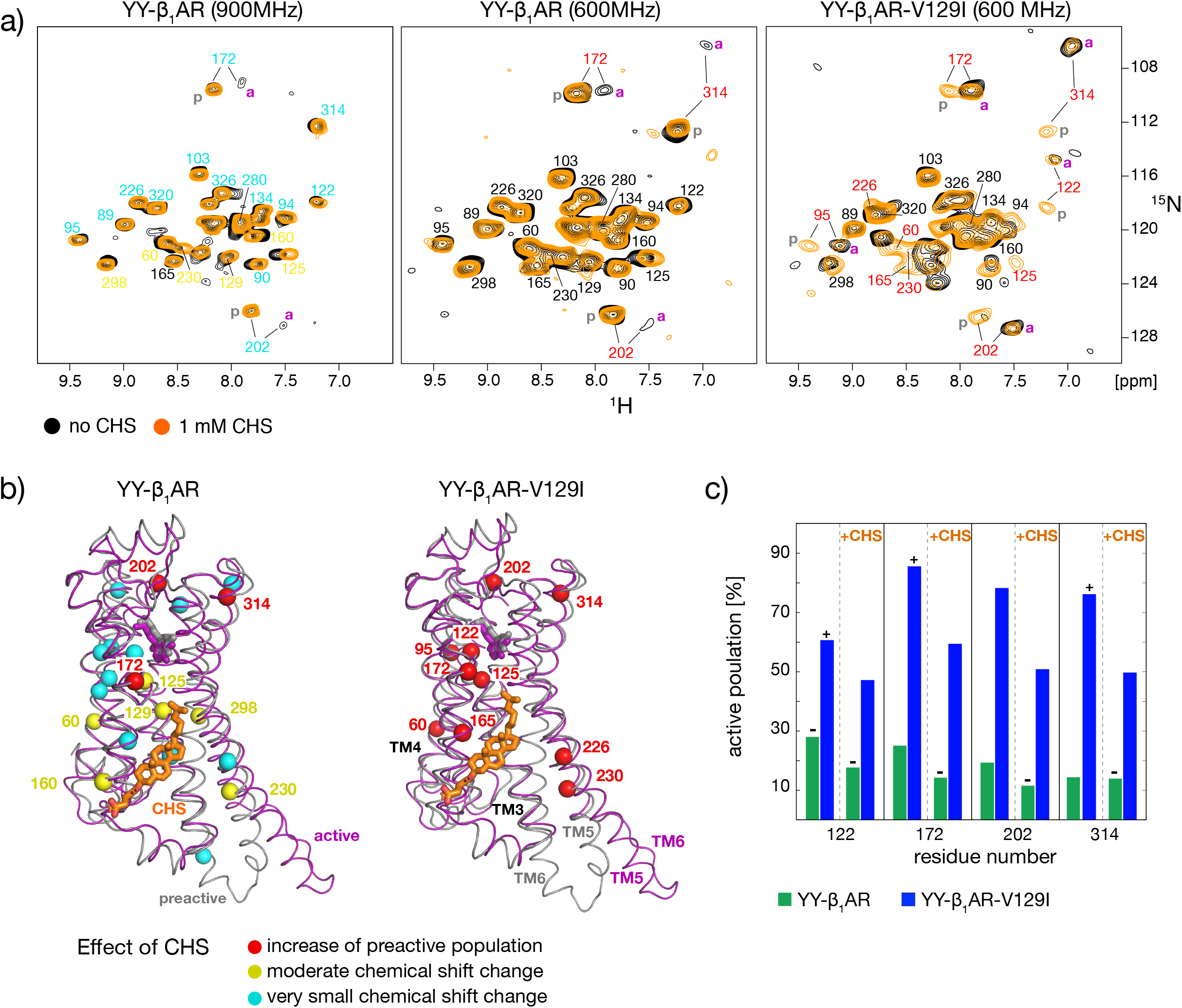
CHS shifts β_1_AR from the active to the preactive conformation. **a)** Superposition of ^1^H-^15^N TROSY spectra of ^15^N-valine-labeled YY-β_1_AR with valine (YY-β_1_AR) or isoleucine (YY-β_1_AR-V129I) at position 129^3.40^ in the absence (black) or presence (orange) of 1 mM CHS. The valine resonances are marked with assignment information using the same color code as in panel **b). b)** Valine residues represented as spheres in the crystal structures (PDB 2y03 - preactive, gray; PDB 6h7j - active, magenta) and colored according to their response to the addition of CHS (red: increase of the preactive conformation population; yellow: moderate chemical shift change; cyan: very small chemical shift change) for YY-β_1_AR (left) and YY-β_1_AR-V129I (right). **c)** Fraction of active-state population of YY-β_1_AR (green) and YY-β_1_AR-V129I (blue) in the absence and presence of CHS. Lower (+) and upper bounds (-) are indicated for cases where only one of the two resonances for the active or preactive conformation could be detected. In these cases, the bounds were derived from the assumption that the amplitude of the missing peak was smaller than three times the root-mean-square deviation of the spectral noise.

### CHS acts as an allosteric inhibitor for β_1_AR

The blocking of the isoprenaline·YY-β_1_AR active state by CHS is already apparent from a careful comparison of ^1^H-^15^N TROSY spectra at 1 bar in the presence and absence of CHS (Figure 3A and S4). Moderate chemical shift perturbations in the vicinity of the CHS binding site detected in the crystal (Figure 3A,B) prove that CHS binds to the same location in the detergent micelle and indicate minor effects on the local conformations of β_1_AR by CHS. As observed previously^5,7^, the resonances of V172^4.56^, V202^ECL2^, and V314^6.59^ are split without CHS into a major and a minor form corresponding to the preactive and active conformation in slow (>5 ms) chemical exchange. Their average intensity ratio corresponds to an active population of ∼19% (Figure 3C and S4). In the presence of CHS, the resonances of the active conformation are no longer visible, indicating that its population must be smaller than ∼13% using the spectral noise as detection limit. Thus CHS shifts the preactive-active equilibrium towards the preactive conformation for isoprenaline·YY-β_1_AR at ambient pressure.

This observation is corroborated by the analysis of a further β_1_AR mutant, YY-β_1_AR-V129I, in which the thermostabilizing mutation I129^3.40^V was reverted to the native isoleucine. In the isoprenaline·YY-β_1_AR-V129^3.40^I complex, the preactive-active equilibrium is strongly shifted towards the active conformation (Grahl et al. in preparation). In the absence of CHS, the ^1^H-^15^N TROSY spectrum of ^15^N-valine-labeled YY-β_1_AR-V129^3.40^I in complex with isoprenaline (Figure 3A) shows very strong active-conformation resonances for V122^3.33^, V172^4.56^, V202^ECL2^, and V314^6.59^ and almost no preactive-conformation resonances. In contrast in the presence of CHS, the resonances of both the preactive and active conformation have very similar intensities. A quantitative analysis of the resonance intensities (Figure 3C) indicates a population of the active conformation larger than ∼80% without CHS, whereas this population decreases to ∼50% in the presence of CHS. Thus, CHS also shifts the preactive-active equilibrium towards the preactive conformation for the YY-β_1_AR-V129^3.40^I mutant in complex with isoprenaline.

Increasing the CHS concentration from 1 to 2 mM further decreased the population of the active conformation for isoprenaline·YY-β_1_AR-V129^3.40^I to ∼40% (Figure S4) showing that the apparent affinity of the CHS binding site must be in the micro-to millimolar range. In contrast, doubling the concentration of isoprenaline from 2 to 4 mM did not significantly change the preactive-active equilibrium (Figure S4). This shows that the orthosteric binding site is fully occupied by isoprenaline and argues against the possibility that CHS competes with isoprenaline for the orthosteric site as suggested previously^59^. Overall, the effects of CHS both on the pressure-dependent β_1_AR spectra and on the preactive-active equilibrium at ambient pressure show that CHS apparently acts as an allosteric inhibitor of the internal activating motions of the receptor.

### Structural basis of β_1_AR regulation by CHS

To rationalize the allosteric inhibition mechanism by CHS, we compared the available crystallographic structures of the binary isoprenaline·β_1_AR complex in the preactive conformation and the ternary complex of isoprenaline·β_1_AR with the G protein-mimicking nanobody Nb80 in the active conformation (Figure 4A). Although CHS was added during the preparation of both complexes^55,60^, an ordered CHS molecule is only detected in the binary preactive complex. In the β_1_AR preactive conformation, CHS makes extensive, hydrophobic contacts to residues in TM3, TM4 and TM5 (Figure 4A and S2C). In the active state, the intracellular part of TM5 swings out of the helical bundle, but the center and extracellular parts move inward and slide by ∼2.5 Å towards the intracellular side (Figure 4A). This would lead to a sterical clash of the I214^5.45^ and I218^5.49^ side chains with the aliphatic end of CHS in the preactive structure. Similarly, the intracellular part of TM4 in the active structure moves towards the CHS position by 1.5 Å and would cause a clash of V160^4.44^ and T164^4.48^ with the hydrophilic tail of CHS. These clashes of CHS with the active β_1_AR conformation may explain the absence of an ordered CHS molecule in the active structure. Besides such repulsive forces, the inactive conformation seems further stabilized by attractive interactions from CHS to residues E130^3.41^, I137^3.48^, and P219^5.50^, which may prevent the inward motion of TM3 and the sliding of TM5 required to reach the active conformation.

**Figure 4.**
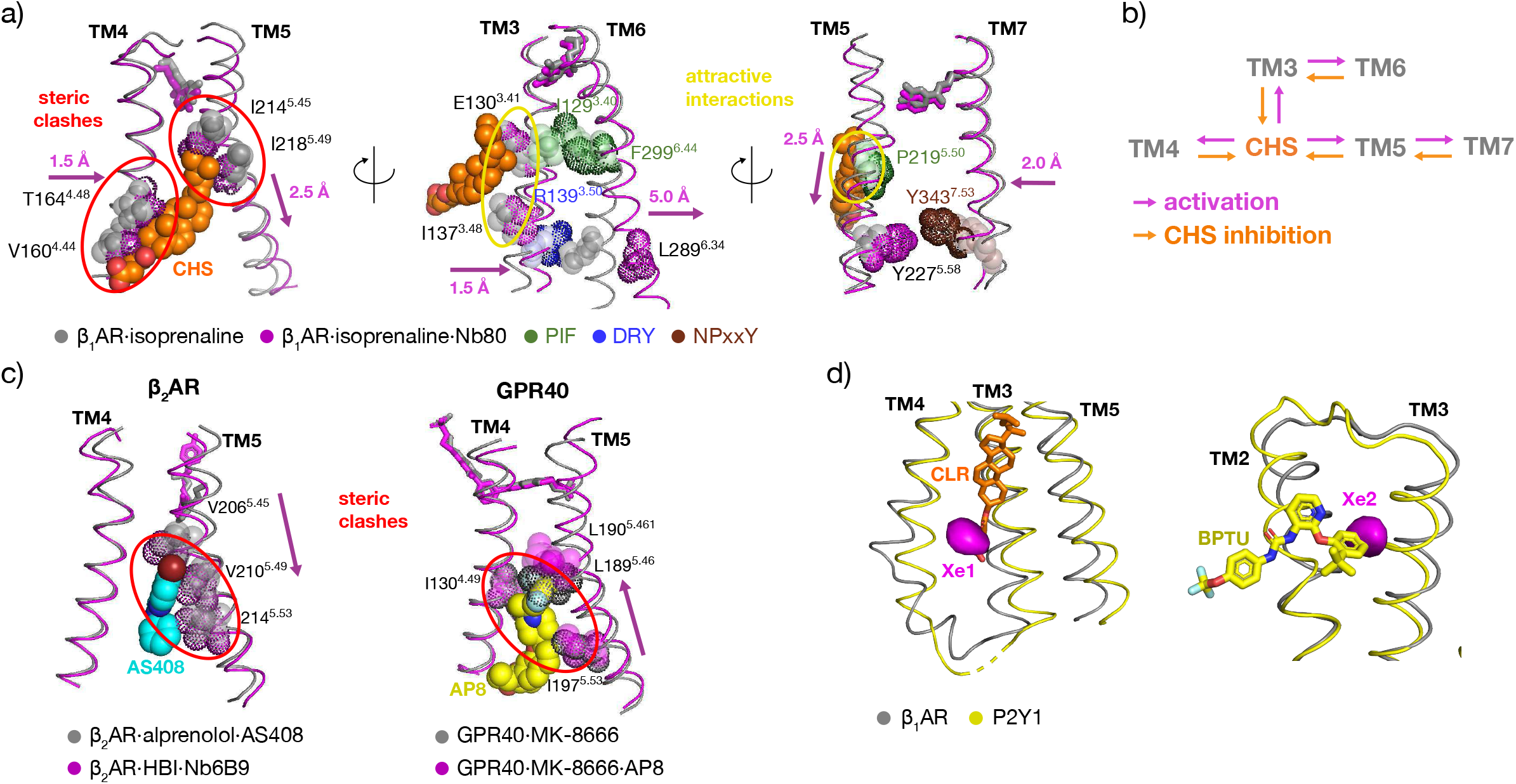
Structural basis of the allosteric inhibition of β_1_AR by CHS. **a)** Comparison between preactive (PDB 2y03, gray) and active (PDB 6h7j, magenta) crystal structures. For clarity only two TMs are shown in each panel together with their key interactions with CHS. Regions of steric clashes of CHS with the active conformation and attractive interactions with the inactive conformation are highlighted. Residues belonging to conserved GPCR microswitch motifs and affected by CHS are shown in varying colors (PIF: green, DRY: blue, NPxxY: brown). The direction of the activating motion is represented by magenta arrows. **b)** Schematic representation of the CHS-transmembrane helix interactions and direction of activating (magenta arrows) and inhibiting (orange arrows, induced by CHS) motions. **c)** Left: comparison of β_2_AR structures in the active (PDB 4ldl, magenta) and preactive state (PDB 6oba, gray) with the negative modulator AS408 (cyan spheres). Right: comparison of agonist-bound MK-8666·GPR40 structures in the absence (PDB 5tzr, gray) and in the presence (PDB 5tzy, magenta) of the positive modulator AP8 (yellow spheres). For both receptors, only the parts of TM4 and TM5 close to the Xe1 binding site are shown together with key residues (spheres) interacting with the allosteric regulators. Receptor residues clashing with the ligand are indicated as dotted spheres. AS408 stabilizes the preactive conformation of TM5 in β_2_AR, but would clash with its active conformation. In contrast, AP8 binding is incompatible with the preactive conformation of GPR40. As a result, the two allosteric modulators shift TM5 in the two receptors in opposite directions (magenta arrows indicate the direction of the activating TM5 motion, see text). **d)** Superposition of the isoprenaline·β_1_AR (PDB 2y03, gray) and the BPTU·P2Y1R (PDB 4xnv, yellow) crystal structures in the vicinity of the β_1_AR Xe1 and Xe2 sites (magenta surfaces). The allosteric antagonist BPTU (yellow) and CLR (orange) observed in the BPTU·P2Y1R structure are shown as sticks.

Taken together, CHS apparently blocks the motions of TM3-5, which activate the canonical GPCR microswitch signaling network (Figure 4A,B). In particular, these comprise the inward motion of residues I129^3.40^ (PIF motif) and R139^3.50^ (DRY motif) in TM3, which push TM6 outwards via F299^6.44^ and L289^6.34^, as well as the downward motion of Y227^5.58^ in TM5, which enables formation of the conserved YY-lock with Y343^7.53^ (NPxxY motif) in TM7 that stabilizes the swung-out position of TM6^7^.

### The detected Xe sites are conserved allosteric binding pockets

To further understand the CHS/CLR allosteric mechanism in the context of other GPCRs, we compared our findings with other receptors, which bind synthetic allosteric modulators at the same Xe1 site (Figure 4C,D). The compound AS408 is a negative allosteric regulator (NAM) of β_2_AR for both G protein activation and arrestin recruitment^61^. The allosteric inhibition of β_2_AR by AS408 can be explained by the same mechanism as the CHS inhibition of β_1_AR, i.e. AS408 appears to hinder the sliding of TM5 towards the intracellular side and to block the TM3 movement, which are necessary for activation. Although the amino acid sequences of β_2_AR and β_1_AR are very similar, including the region of this allosteric binding site, the substitution of a single amino acid at position 3.41 (valine in β_2_AR and leucine in β_1_AR) is sufficient to reduce the affinity of AS408 for β_1_AR tenfold relative to β_2_AR^61^. This substitution may also be the reason why β_2_AR does not bind cholesterol at the same site.

A further comparison with the receptor GPR40 (free fatty acid receptor 1) is particularly interesting, since the binding of the allosteric compound AP8 to the same site as occupied by CHS in β_1_AR induces an upregulation of activity. Thus AP8 is a positive allosteric modulator (PAM) enhancing the response of partial agonists^62^. No active structures of GPR40 are available in complexes with a G protein or G mimic. However, TM5 of the preactive GPR40 in complex with the agonist MK-8666 is shifted towards the intracellular side relative to other active GPCR·G protein complexes (β_2_AR, A2R)^62^. Binding of the PAM AP8 to the MK-8666·GPR40 complex then shifts TM5 towards the extracellular side (Figure 4C), i.e. towards the conformation of the active β_2_AR and A2R GPCR complexes. Thus the shift of TM5 from the preactive to the active GPR40 conformation is opposite to the usual direction. The alignment of the binary MK-8666·GPR40 and ternary MK-8666·GPR40·AP8 complexes (Figure 4C) reveals that AP8 would clash with the preactive conformation of TM5, thereby explaining its positive modulation of GPR40 activity, in complete agreement with the negative modulation by CHS and AS408 binding to the same site in β_1_AR and β_2_AR, respectively.

A further structure, i.e. P2Y1R in complex with the inhibiting antithrombotic drug BPTU (PDB 4XNV)^63^ also has a CLR molecule bound close to the Xe1 site in β_1_AR albeit in a different orientation (Figure 4D). The effect of CLR on P2Y1R has not been characterized in detail, but P2Y1R-mediated calcium release in platelets was reported as insensitive to CLR depletion^64^. Intriguingly, however, the antithrombotic drug BPTU binds to a second allosteric pocket on the external P2Y1R interface with the lipid bilayer (Figure 4D), which coincides with the Xe2 site in β_1_AR. Thus apparently, albeit detected in a different receptor, also the Xe2 site constitutes a drug binding pocket.

## DISCUSSION

Here we have determined and rationalized in atomic detail the effect of CHS onto the β_1_AR functional equilibria by a combination of crystallography and solution NMR. As high pressure shifts the preactive-active conformational equilibrium of the isoprenaline·YY-β_1_AR complex to the active conformation, the receptor must compress empty, hydrophobic cavities during activation. We have localized such cavities by X-ray diffraction of Xe-derivatized β_1_AR crystals. Intriguingly, one of the detected cavities colocalizes with the CHS binding site. Our NMR analysis shows that CHS abrogates the pressure-induced switch of the isoprenaline·YY-β_1_AR complex to the active conformation and shifts the preactive-active equilibrium to the preactive conformation. Therefore CHS acts as an allosteric inhibitor of β_1_AR.

The inhibitory effect of CHS can be rationalized by a comparison of the preactive isoprenaline·β_1_AR structure with bound CHS and the active isoprenaline·β_1_AR·Nb80 structure, which does not contain detectable CHS. In the binary isoprenaline·β_1_AR complex, CHS stabilizes the preactive conformation by numerous hydrophobic interactions. However, it would clash with the active conformation. Thereby CHS obstructs functional motions required for the activation of the canonical GPCR microswitch network.

As compared to CHS, even stronger inhibitory effects are expected for CLR due to its larger hydrophobicity and the hydrophobic character of the identified combined Xe- and CHS-binding site. This inhibitory function of CLR is corroborated by the enhanced signaling of α- and β-adrenergic receptors in cardiomyocytes depleted of cellular CLR^65^. Being highly abundant in the cell membrane [34% of the total lipid content in mammalian plasma membranes^33,37^], CLR may be a prime regulator of adrenergic receptors, keeping their basal activity low by stabilizing their preactive conformation. Very likely, the activity of many further GPCR is modulated by CLR. As CLR levels vary among different cell types and subcellular compartments, this may introduce an additional layer of cell-specific receptor modulation.

The identified CLR binding sites in GPCRs are very diverse. Therefore there cannot be a universal mechanism of allosteric GPCR regulation by CLR. However, our study reveals two general aspects of the CLR action on GPCRs: (i) by filling dry cavities, CLR reduces the total volume available for functional motions and (ii) CLR ties together hydrophobic parts of the receptor thereby stabilizing a particular conformation.

Several allosteric modulators of other GPCRs bind to the same site as CHS in β_1_AR. Although their binding site is identical, the functional output may vary. For example, such allosteric modulators increase the activity of GPR40 and DRD1^62,66^, but down-modulate β_2_AR and C5aR1^61,67^. Blocking TM5 sliding appears as the hallmark of their mechanism of action with the position of the blocked TM5 relative to the active conformation defining the direction of modulation.

The present study adds to the mounting evidence that empty cavities may fulfill key functions in proteins such as the high-affinity binding of hydrophobic ligand moieties^2^ or directing functional motions^5^. Apparently, CHS fills a void that must be compressed for β_1_AR activation. Filling spaces between the transmembrane helices modulates the dynamics and activity of many GPCRs, as demonstrated by the development of a number of positive and negative allosteric modulators^61– 63,66–69^, which occupy such crevices. In contrast to the orthosteric binding sites, the surfaces of allosteric binding sites are often lined by amino acids that are distinctive between GPCR subtypes. This property gives them a high potential for the development of therapeutic drugs targeting specific GPCR subtypes.

Intriguingly, not only the detected xenon binding site Xe1 in β_1_AR colocalizes with the binding site of the allosteric modulator CHS/CLR, but also Xe2 colocalizes with the antithrombotic drug BPTU in P2Y1R. Therefore the detection of empty cavities by xenon may present a valuable method to screen for new drug target sites in GPCRs.

## METHODS

### Protein expression and purification

Expression of the unlabeled, thermostabilized turkey β_1_AR construct (TS-β_1_AR), as well as of ^15^N-valine-labeled G protein binding-competent construct (YY-β_1_AR) in baculovirus-infected Sf9 cells, purification, assignments, binding of ligands, and exchange between ligands were carried out as described previously^8^. The plasmid for Nb80 was a generous gift by Jan Steyaert and the Nb80 protein was purified according to the described procedure^70^.

For the TS-β_1_AR crystallization sample, an additional size-exclusion purification step was added to increase the sample homogeneity and to exchange the detergent from DM to Hega-10. For this, ∼900 μL of 100 μM TS-β_1_AR was applied at 0.3 mL/min to a 24-mL Superdex 200 Increase 10/300 GL column (GE healthcare) pre-equilibrated with 10 mM Tris-HCl, 100 mM NaCl, 0.1 mM EDTA, 1 mM isoprenaline, 0.35% Hega-10, pH 7.5. The receptor was eluted at 0.3 mL/min with the same buffer, and concentrated with a 50-kDa molecular weight cutoff centrifugal filter (Amicon) to a final concentration of ∼10– 15 mg/mL.

### Crystallization

Immediately before setting up the crystallization plates, CHS was added to the isoprenaline·TS-β_1_AR complex to a final concentration of 0.45 mg/mL (from a 10 mg/mL stock solution in 2% Hega-10). The receptor solution was subsequently centrifuged at 130’000 g through a 0.22-μm Ultrafree-MC spin filter (Millipore) for 5 minutes at 4°C in order to remove possible aggregates. Crystals were grown under similar conditions as described previously^55,71^ on MRC Maxi 48-well crystallization plates (Swissci) by vapor diffusion in sitting drops containing 1 μL of protein solution + 1 μL of reservoir solutions consisting of 0.1 M Tris-HCl pH 7.5–9.0, and 20–30% PEG 600. Plates were incubated at 4°C until crystals reached their maximum size after ∼2 weeks. Typically, the obtained crystals were rod-shaped with dimensions of ∼250×75×30 μm^3^.

### Xenon derivatization

Xenon derivatization of the isoprenaline·TS-β_1_AR crystals was not possible with the commonly used commercial Xcell (Oxford Cryosystems), since the fragile GPCR crystals dried and broke after a few minutes of pressurization. For this reason, we built a pressure cell equipped with a wash bottle for humidity equilibration as prechamber and a humidity sensor, in which the humidity can be adjusted to arbitrary values using suitable aqueous salt solutions in the wash bottle. A relative humidity of 100% produced by pure water proved suitable to maintain TS-β_1_AR crystals stable for long periods without changes in their appearance. After the xenon incubation, the xenon pressure was released and the crystals were flash-frozen in liquid nitrogen within seconds. Derivatization conditions were varied between 5–30 min incubation and 5–12 bar xenon pressure. The strongest anomalous signal was obtained for crystals incubated for 20 minutes at a xenon pressure of 5 bar. Cryo-protectant soaking with PEG 600 before crystal incubation with xenon did not improve crystal quality.

### Data collection and structure determination

X-ray diffraction data were collected at the Swiss Light Source, Paul Scherrer Institute (Villigen, Switzerland), using the PXIII beamline equipped with a PILATUS 2MF detector (Dectris, Baden-Dättwil, Switzerland). Isoprenaline·TS-β_1_AR crystals belonged to the space group *P*2_1_ with the following unit cell parameters a=59.324 Å, b=125.653 Å, c=87.774 Å and β=104.76°. The crystal structure was solved by molecular replacement using the atomic coordinates of the β_1_AR from PDB entry 2y03 (chain A) as a template, and searching for two β_1_AR molecules using the program PHASER^72^ contained in the CCP4i2 (version 1.0.2) package^73^. In order to maximize the anomalous contribution of the Xe atoms, diffraction data were recorded at 6.0 keV on four different crystals under varying *chi* angles. The individual *chi-*angle data sets were then processed using the autoPROC pipeline^74^ and merged for each crystal with XSCALE^75^. Identification of Xe atoms was performed by computing Fourier maps using SHELXC^76^ and ANODE^77^. The two xenon binding sites (Xe1 and Xe2) described in the manuscript correspond to the strongest anomalous signals detected in all four crystals and present in both chains (A and B) of the asymmetric unit (Table S2).

### Analysis of crystal structures

All molecular representations were generated using the PyMOL 2.1. Molecular Graphics System^78^. The various receptor structures were aligned on the following TM regions: 1.46–1.51, 2.46–2.56, 3.34–3.44, 4.48–4.56, 7.38–7.46. Voids in the crystal structure of isoprenaline·β_1_AR (PDB 2y03) were identified and quantified using the program Hollow 1.2.^79^ as described previously^5^ using a grid spacing of 1.0 Å and a sphere size of 4 Å for defining the receptor surface.

### NMR experiments

NMR samples were prepared with typical receptor concentrations of 100 −200 μM in 20 mM Tris-HCl, 100 mM NaCl, ∼40 mM decylmaltoside (DM), 0.02% NaN_3_, 5% D_2_O (10% for high-pressure experiments), pH 7.5. The isoprenaline·β_1_AR complex was formed by adding 2 mM isoprenaline and 20 mM sodium ascorbate (to prevent oxidation of the ligand) to the apo receptor. For formation of the ternary complex, a 1.2-molar equivalent of Nb80 was added to the isoprenaline·β_1_AR complex. For measurements in the presence of 1 mM or 2 mM CHS, suitable volumes of 10 mg/mL CHS stock solution in 20 mM Tris-HCl pH 7.5, 100 mM NaCl, 0.1 % DM were added directly to the NMR sample.

The ^1^H-^15^N TROSY NMR experiments were recorded on Bruker AVANCE 900 MHz or AVANCE 600 MHz spectrometers equipped with TCI cryoprobes at 304 K using sample volumes of ∼250 μL either in Shigemi microtubes or a commercial high-pressure NMR cell (3 mm inner diameter, 120 μl active volume, rated to 2,500 bar, Daedalus Innovations LLC) as described previously^5^. The sample stability during longer NMR experiments was monitored by interleaved, one-dimensional ^1^H experiments with spin-echo water suppression^80^.

## DATA AVAILABILITY

Structure factors, phases and density maps derived from the anomalous scattering data of the four xenon-derivatized isoprenaline·β_1_AR crystals as well the β_1_AR structure derived from the first crystal have been deposited in the Zenodo repository under DOI 10.5281/zenodo.4926013.

## ACKNOWLEDGMENTS

This work was supported by the Swiss National Science Foundation (Grants CRSK-3_195592 to L.A.A., and 31-149927, 31-173089, 31-201270 to S.G.). We gratefully acknowledge Michel Schaffhauser, Patrick Schlenker and Raymond Strittmatter (Biozentrum Central Mechanical Workshop) as well as Simon Saner (Biozentrum Central Electronics Workshop) for designing and building the xenon pressure chamber apparatus, Dr. Timothy Sharpe (Biozentrum Biophysics Facility) for expert help with the biophysical characterization of β_1_AR, the Paul Scherrer Institut, Villigen, Switzerland for provision of synchrotron radiation beamtime at beamline PXIII, as well as Drs. Hans-Jürgen Sass, Timm Maier, and Tilman Schirmer for helpful discussions.

## AUTHOR CONTRIBUTIONS

L.A.A. and S.G. conceived the study. L.A.A. and A.G. expressed and purified protein and recorded NMR spectra. L.A.A. recorded high-pressure NMR experiments. L.A.A. analyzed and interpreted the NMR data. R.D.T. and S.E. recorded the X-ray data. R.D.T. analyzed the X-ray data. L.A.A., R.D.T. and S.G. wrote the manuscript.

## COMPETING INTERESTS STATEMENT

The authors declare no competing interests.

## Notes

### Competing Interest Statement

The authors have declared no competing interest.

