## Supplementary material for "Filling of a water-free void explains the allosteric regulation of the β_1_-adrenergic receptor by cholesterol"

### Supporting Information

*Address correspondence to:
Layara Abiko
Focal Area Structural Biology and Biophysics, Biozentrum
University of Basel, CH-4056 Basel, Switzerland
Phone: ++41 61 267 2100
FAX: ++41 61 267 2109


Stephan Grzesiek
Focal Area Structural Biology and Biophysics, Biozentrum
University of Basel, CH-4056 Basel, Switzerland
Phone: ++41 61 267 2100
FAX: ++41 61 267 2109


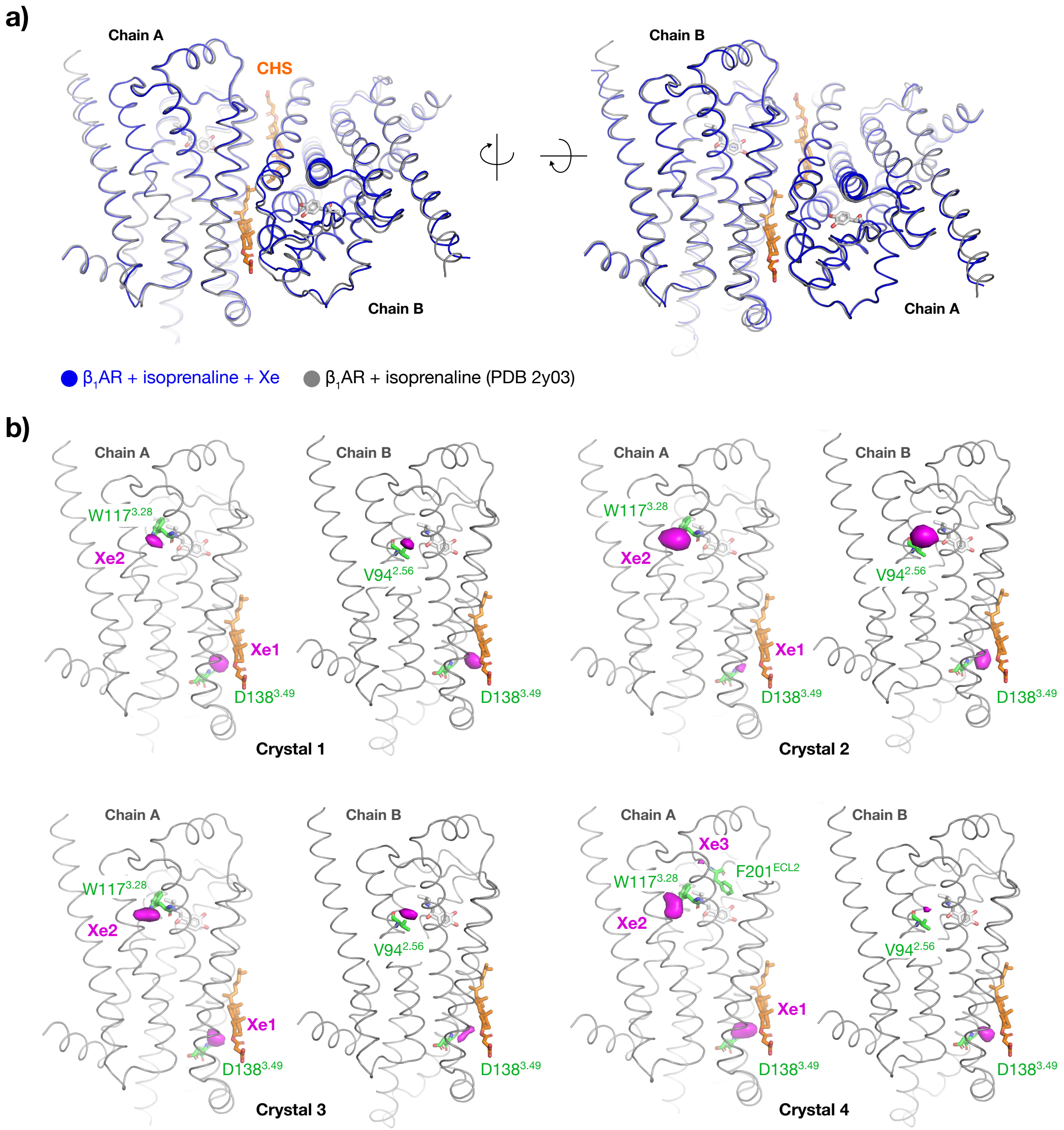


**Figure S1. X-ray analysis of Xe-derivatized isoprenaline·TS-β$_1$AR crystals. a)** Superposition of the structure determined from the Xe-derivatized isoprenaline·TS-β$_1$AR crystals and the structure solved in the absence of xenon (PDB 2y03). **b)** Position of the Xe anomalous scattering density (magenta) detected in the four different crystals superimposed onto the structure of the isoprenaline·β$_1$AR complex (PDB 2y03). Anomalous scattering signals with a height larger than 5σ are marked by the numbering (Xe1–Xe3) given in Table S2. For each xenon site, the nearest amino acid (determined from the distance to the backbone nitrogen atom) is shown as green sticks. Across all crystals, sites Xe1 and Xe2 are consistently located at the same position within the two β$_1$AR chains of the asymmetric unit.

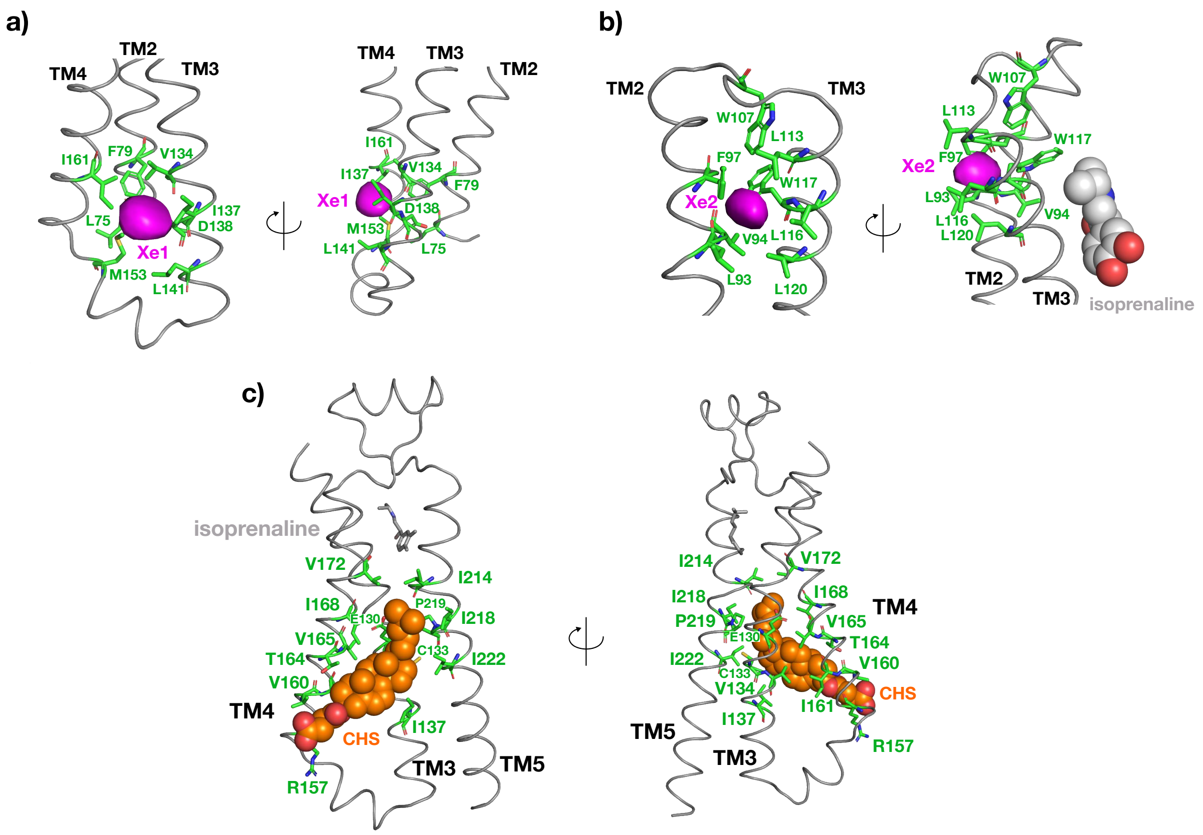


**Figure S2. Hydrophobic interactions of xenon and CHS with the β$_1$AR side chains.** Close-up of the detected **a)** Xe1 and **b)** Xe2 xenon sites (magenta) superimposed onto the PDB 2y03 crystal structure. Side chains forming the hydrophobic Xe binding sites are represented by green sticks and isoprenaline as gray spheres, respectively. **c)** Hydrophobic interactions between CHS (orange spheres) and β$_1$AR side chains (green sticks).

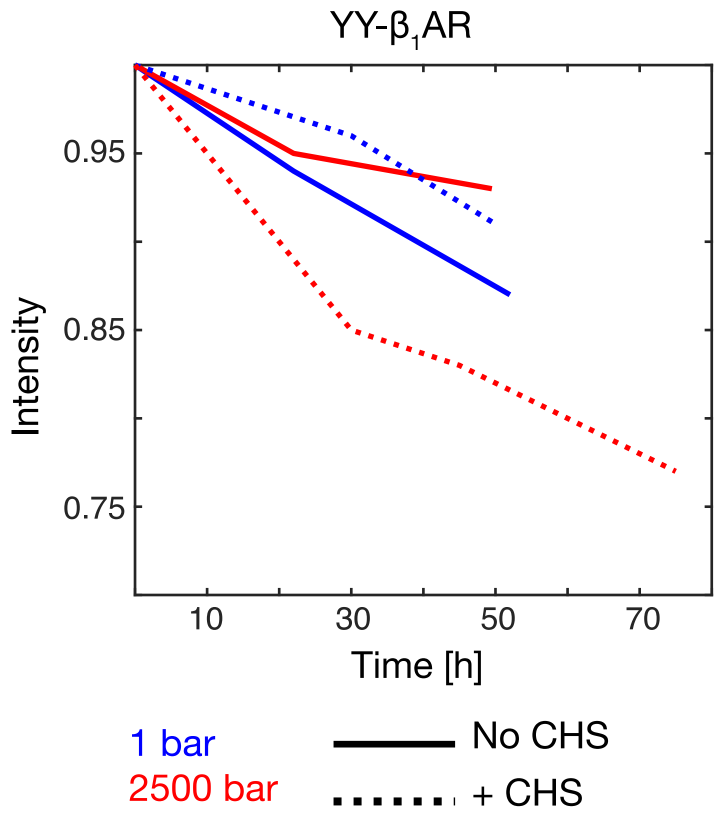


**Figure S3. Stability of YY-β$_1$AR sample in the NMR tube at 304 K.** The normalized peak intensity of amide protons (8.3 ppm) is shown over time for YY-β$_1$AR in the absence (solid lines) or presence (dashed lines) of 1 mM CHS at 1 bar (blue) or 2500 bar (red).

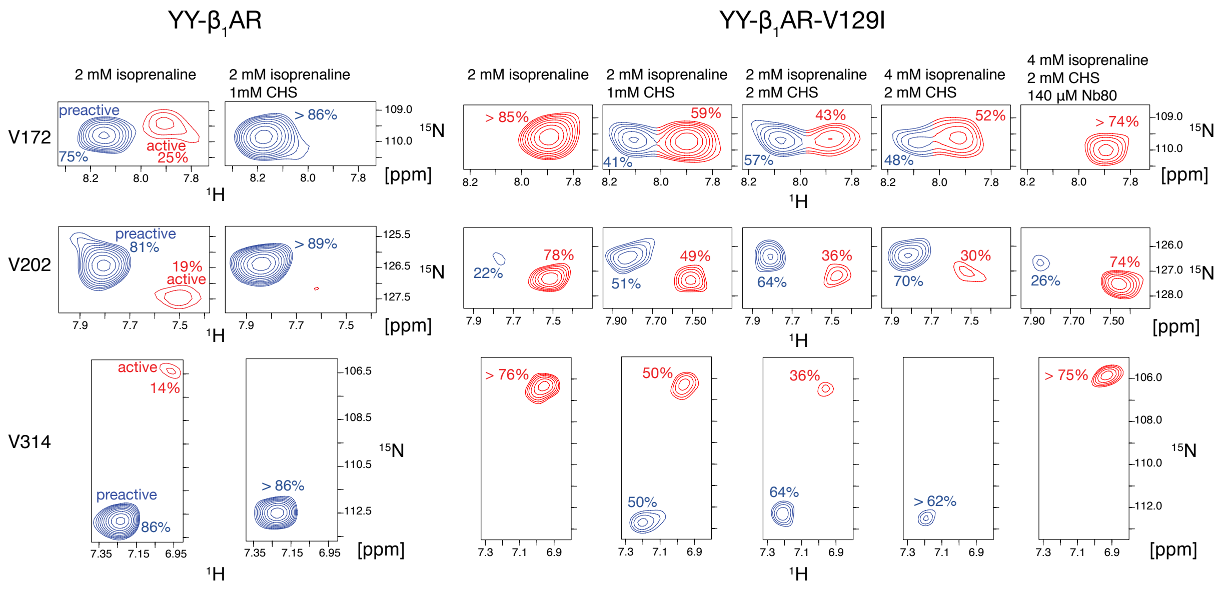


**Figure S4. Effect of CHS, isoprenaline and Nb80 on the β$_1$AR preactive-active equilibrium.** Zoomed regions for resonances of V172$^{4.56}$, V202 $^{ECL2}$, and V314$^{6.59}$ in the $^{1}$H-$^{15}$N TROSY spectra of the isoprenaline·YY-β$_1$AR and isoprenaline·YY-β$_1$AR-V129$^{3.40}$I complexes are shown under varying concentrations of isoprenaline, CHS and Nb80. Resonances of the preactive and active state are coloured in blue and red, respectively. The population fractions of the preactive and active state derived from the resonance intensities are indicated.

**Table S1.** Data collection and refinement statistics for the $\beta_1$AR structure derived from the xenon-derivatized isoprenaline·TS-$\beta_1$AR crystal.

| | |
|---|---|
| **Synchrotron source** | SLS, PXIII |
| **Resolution [Å]** | 3.4 |
| **Space group** | $P2_1$ |
| **a, b, c [Å]** | 59.324, 125.653, 87.774 |
| **α, β, γ [°]** | 90.00, 104.76, 90.00 |
| **Total reflections** | 64667 (2801) |
| **Unique reflections** | 9550 (478) |
| **Multiplicity** | 6.8 (5.9) |
| **Completeness [%]** | 89.8 (56.9) |
| **Mean I/sigma(I)** | 11.3 (1.1) |
| **R-merge** | 0.054 (1.890) |
| **R-meas** | 0.059 (2.074) |
| **CC1/2** | 0.999 (0.573) |
| **Anomalous multiplicity** | 3.5 (3.1) |
| **Anomalous completeness [%]** | 89.2 (55.0) |

Statistics for the highest-resolution shell are shown in parentheses.

**Table S2.** Position and height of strongest anomalous Xe scattering signals (>5σ) detected in four different xenon-derivatized isoprenaline·$\beta_1$AR crystals.

**Crystal 1**

| Site[a] | X | Y | Z | Height [σ] | Nearest residue[b] |
|---|---|---|---|---|---|
| **Xe1_A** | -6.677 | 4.500 | 12.546 | 9.09 | $D138^{3.49}$ |
| **Xe1_B** | -6.358 | -5.955 | 40.302 | 7.72 | $D138^{3.49}$ |
| **Xe2_A** | -24.320 | -19.526 | 15.122 | 6.19 | $W117^{3.28}$ |
| **Xe2_B** | 13.070 | -21.573 | 23.626 | 6.07 | $V94^{2.56}$ |

**Crystal 2**

| Site | X | Y | Z | Height [σ] | Nearest residue |
|---|---|---|---|---|---|
| **Xe1_A** | -6.584 | 5.270 | 11.237 | 4.63 | $D138^{3.49}$ |
| **Xe1_B** | -5.253 | -5.677 | 41.483 | 5.40 | $D138^{3.49}$ |
| **Xe2_A** | -25.051 | -19.382 | 14.798 | 7.58 | $W117^{3.28}$ |
| **Xe2_B** | 13.175 | -23.393 | 22.673 | 7.27 | $V94^{2.56}$ |

**Crystal 3**

| Site | X | Y | Z | Height [σ] | Nearest residue |
|---|---|---|---|---|---|
| **Xe1_A** | -7.052 | 5.299 | 12.570 | 6.52 | $D138^{3.49}$ |
| **Xe1_B** | -5.443 | -5.643 | 40.028 | 5.19 | $D138^{3.49}$ |
| **Xe2_A** | -24.930 | -18.147 | 16.113 | 6.55 | $W117^{3.28}$ |
| **Xe2_B** | 13.176 | -21.926 | 23.219 | 5.69 | $V94^{2.56}$ |

**Crystal 4**

| Site | X | Y | Z | Height [σ] | Nearest residue |
|---|---|---|---|---|---|
| **Xe1_A** | -6.234 | 4.378 | 12.275 | 7.19 | $D138^{3.49}$ |
| **Xe1_B** | -4.870 | -5.565 | 41.331 | 6.76 | $D138^{3.49}$ |
| **Xe2_A** | -25.183 | -18.241 | 14.474 | 6.63 | $V94^{2.56}$ |
| **Xe2_B** | 13.578 | -23.096 | 22.767 | 5.00 | $V94^{2.56}$ |
| **Xe3_A** | -25.475 | -21.107 | 26.662 | 6.46 | $F201^{ECL2}$ |

[a]A or B indicate the $\beta_1$AR chain in PDB 2y03.

[b]Amino acid residue nearest to the xenon site according to the backbone nitrogen position (PDB 2y03).
